# *Bacteroides* vesicles promote functional alterations in gut microbiota composition

**DOI:** 10.1101/2024.03.05.583531

**Authors:** Olga Yu. Shagaleeva, Daria A. Kashatnikova, Dmitry A. Kardonsky, Boris A. Efimov, Viktor A. Ivanov, Suleiman S. Evsiev, Eugene A. Zubkov, Olga V. Abramova, Yana A. Zorkina, Anna Y. Morozova, Elizaveta A. Vorobeva, Artemiy S. Silantiev, Irina V. Kolesnikova, Maria I. Markelova, Evgenii I. Olekhnovich, Maxim D. Morozov, Polina Y. Zoruk, Daria I. Boldyreva, Anna A. Vanyushkina, Andrei V. Chaplin, Tatiana V. Grigoryeva, Natalya B. Zakharzhevskaya

## Abstract

Inflammatory bowel diseases are characterized by chronic intestinal inflammation and alterations of gut microbiota composition. *Bacteroides fragilis*, which secretes outer membrane vesicles (OMVs) with polysaccharide A (PSA), can moderate inflammatory response and possibly alter microbiota composition. In this research, we created a murine model of chronic DSS- induced intestinal colitis and treated it with *Bacteroides fragilis* OMVs. We monitored the efficiency of OMVs therapy by determining the disease activity index (DAI) and histological examination (HE) of the intestine before and after vesicles exposure. We also analyzed the microbiota composition using 16S rRNA gene sequencing. Finally, we evaluated the volatile compounds composition in animals’ stools by HS-GC/MS to assess the functional activity of the microbiota. As a result, we observed a more effective intestine repair after OMVs treatment according to DAI and HE. The metabolomic study also indicated the microbiota functional activity change, showing a predominance of phenol and pentanoic acid in the control group compared to the group treated with DSS (DSS) and the group treated with OMVs (DSS OMV). We also observed a positive correlation of these metabolites with *Saccharibacteria* and *Hungatei Clostridium* in the control group, whereas in the DSS group there was a negative correlation of phenol and pentanoic acid with *Lactococcus* and *Romboutsia*. According to metabolome and sequencing data, the microbiota composition of the DSS OMV group was intermediate between the control and DSS groups. It can be concluded that OMVs not only have an anti-inflammatory effect, but also contribute to the recovery of the microbiota composition.

## Introduction

IBD is a heterogeneous chronic and relapsing inflammatory disease, traditionally divided into ulcerative colitis (UC) and Crohn’s disease (CD) [1–3]. Both UC and CD can cause various symptoms such as diarrhoea, rectal bleeding and abdominal pain. However, inflammation in CD is transmural and can affect any part of the gastrointestinal tract. In contrast, inflammation in UC is more superficial and limited to the colon [4]. Although the pathogenesis of IBD remains unknown, it has been described as a multifactorial disease involving both genetic and environmental components [5]. Changes in the gut microbiota have also been associated with IBD. Standard IBD therapy aims to reduce inflammation with hormones and antibiotics. At the same time, antibiotics cause changes in the microbiota composition, which contributes to increased inflammation [6]. This creates a vicious cycle and provokes the development of chronic inflammation in the gut. Current probiotics are largely ineffective because bacterial colonization occurs during the acute phase of inflammation, when survival conditions are extreme. Bacteria may develop a pathogenic phenotype when its adapting to chronic inflammation in intestine [7]. In this regard, it is important to select microorganisms that are able to suppress inflammation and promote effective colonization.

Among bacteria with immunomodulatory properties, Bacteroides are the best known because of the surface located polysaccharide A (PSA) [8]. PSA interacts with TLR2 receptors on dendritic cells, leading to the synthesis of interleukin-10. Interleukin-10 molecules modulate the activity of CD4+FOXP3+ regulatory T cells (Tregs), which suppress inflammation [9–10]. PSA is also found on the outer membrane vesicles (OMV) surface which are produced in large quantities by *Bacteroides fragilis* [11]. *Bacteroides* species cannot be used as a single component of probiotics because it can change its phenotype very quickly, from non-pathogenic to pathogenic strain [12]. However, the use of the *Bacteroides* OMVs with PSA may be justified. Previously it has been shown that PSA prevents experimental colitis in murine model. [13]. However, the efficiency of OMVs as a therapeutic agent has not been fully evaluated. In our previous research, we performed a detailed proteomic and metabolomic study of OMVs produced by the non-toxigenic *Bacteroides fragilis* strain JIM10 [14]. Combined with HS-GC-MS data, the proteomic and metabolomic results of *Bacteroides fragilis* OMVs analysis showed that vesicles contain several enzymes, metabolites and volatile compounds, especially fatty acids [15]. Multi-component OMVs are expected to be more effective than isolated PSA in reducing inflammation and subsequent microbial colonization [16].

Models of DSS-induced colitis are the most widely used in IBD research and are also optimal for studying therapeutic agents [17–18]. DSS-induced colitis can be characterized by histological analysis and measurement of the disease activity index (DAI) [19]. The DAI includes general characteristics: stool consistency, including the presence of blood, animal behavior and weight. Traditionally, it has been assumed that DSS does not alter the microbiota composition. However, a number of recent studies have suggested that DSS may indirectly affect species diversity [20]. A recent study of the microbiota composition in DSS-induced colitis model identified two species, *Duncaniella muricolitica* and *Alistipes okayasuensis*, that were associated with worse disease outcome after DSS treatment [21]. These pathogenic taxa reproduced a severe DSS response when used to noncolonized germ-free mice, confirming the causal relationship between these species and the severity of DSS colitis. It is therefore important to assess the microbiota composition when DSS is used. However, significant changes in microbiota composition are not expected. Metagenomic research is traditionally used to characterize the microbiota diversity [22]. Volatile metabolites released by bacteria also can be useful for functional activity of the microbiota assessment [23].

In this study, we created a DSS-induced colitis murine model and tested *Bacteroides fragilis* JIM10 strain OMVs as a therapeutic agent for intestine recovery. We evaluated the effectiveness of OMVs therapy by counting DAI and HE based on histology examination for all groups. In addition, we determined the relative amounts of volatile compounds by HS GC-MS and analyzed the composition of the gut microbiota by 16S rRNA sequencing.

## Material and methods

### Bacterial strain and growth conditions

*Bacteroides fragilis* (JIM10 strain - NZ_MBRB00000000.1) was stored lyophilised at -80°C in 20% (w/v) sucrose and 1% (w/v) gelatin solution. The strain was grown anaerobically on Anaerobe Basal Agar (Oxoid) supplemented with 5% (v/v) defibrinated sheep blood under anaerobic conditions established by placing Anaerogen™ bags in anaerobic 3.5 litre flasks (Oxoid; Thermo Fisher Scientific, Inc.) or in anaerobic flasks (Schuett-Biotec, Germany) at 37°C until stationary phase. For liquid culture, a preculture of JIM10 strain was grown anaerobically in Columbia Broth Base (Hi Media, India) at 37°C until stationary phase.

### OMV isolation

Two hundred milliliters of a 24-hour liquid bacterial culture of *Bacteroides fragilis* JIM10 was centrifuged at 4,500g at 4°C. To remove residual cells, the supernatant was filtered through a 0.45 μm porous membrane. (The filtrate was subjected to ultracentrifugation at 100,000g for 2 h (Optima L-90 K ultracentrifuge; Beckman Coulter). The supernatant was discarded and the pellet was washed with sterile PBS and filtered through a sterile 0.2 μm-pore polyvinylidene difluoride (PVDF) membrane (Millex GV; Millipore). Ultracentrifugation was repeated twice. The vesicle pellet was resuspended in distilled water or 150 mM NaCl (pH 6.5). The concentration of OMVs was quantified as the dry precipitate.

### Electron microscopy

Isolated *Bacteroides fragilis* JIM10 OMVs were diluted to a concentration in the range of 2-5 x 10^11 particles/mL and prepared for TEM analysis as previously described [15]. Images were obtained using a JEM-1400 (Jeol, Tokyo, Japan) transmission electron microscope equipped with a Rio-9 camera (Gatan Inc., Pleasanton, CA, USA) at 120 kV.

### Ethics Statement

All experimental procedures were set up and maintained in accordance with Directive 2010/63/EU of 22 September 2010 and approved by the local ethical committee of V.P. Serbsky National Medical Research Center for Psychiatry and Narcology.

### Animals

Female C57BL/6 mice, 2 months old (20-25 g), obtained from the Laboratory Animal Nursery (Pushchino, RAS, Moscow Region) were used in the experiment. The animals were randomly divided into three groups: the control group (n=10), the sodium dextran sulfate (DSS) group (n=10) and the sodium dextran sulfate group treated with outer membrane vesicles (DSS+OMV) (n=10). Each mouse in the DSS group received 4% DSS orally for 5 days. Intestinal inflammation was maintained for 20 days in the DSS group. During this time, animals were alternately given water with and without DSS. All animals were then euthanized with chloroform. All animals in the DSS+OMV group were maintained under the same conditions as the DSS group till the 10^th^ day. From day 10, animals with persistent inflammation were treated with OMVs (1 µg/kg). All mice from were DSS+OMV group were euthanized with chloroform on day 20. The control group of mice received the same volume of normal saline for 20 days prior to euthanasia by chloroform. Body weight, stool consistency and the presence of occult blood were measured every 3 days. Stool samples were collected for subsequent metabolomic and 16S rRNA gene sequencing.

### Reagents

Dextran sulfate sodium salt, Mr ∼40,000, Alfa Assar. N, O- bis(trimethylsilyl)trifluoroacetamide (with 1% trimethylchlorosilane; v/v; lot no. B-023), and heptadecanoic acid (as an internal standard, IS, purity ≥ 98%; lot no.H3500) were purchased from Sigma-Aldrich (Saint Louis, USA). O-methyl hydroxylamine hydrochloride (purity: 98.0%; lot no. 542171) was purchased from J&K Scientific Ltd. (Beijing, China). Pyridine (lot no. C10486013) was obtained from Macklin Biochemical (Shanghai, China). Chromatographic-grade methanol was purchased from Thermo Fisher Scientific (Waltham, USA). Pure water was obtained from Wahaha Company (Hangzhou, China).

### Clinical Disease Score

The Disease Activity Index (DAI) was estimated by the score of body weight loss (no weight loss: 0; 5-10% weight loss: 1; 11-15% weight loss: 2; 16-20% weight loss: 3; 20% weight loss: 4), stool consistency (formed: 0; watery stool: 2) and the degree of stool occult blood (normal stool color: 0; reddish stool color: 2).

### Histology

Colon samples were cut lengthwise, washed with PBS and rolled into a “Swiss roll” 20. The obtained material was fixed in 10% buffered formalin pH 7.0-7.8 for 48 hours and then placed in labeled histological cassettes with a liner to prevent unrolling. Histological processing was performed with isopropyl alcohol (Chimmed, Russia) and two changes of o-xylene (Chimmed, Russia). The processed samples were embedded in paraffin (BioVitrum, Russia). Sections of 3.5 μm thickness were cut on a Thermo HM 340E rotary microtome (Thermo Scientific, China, manufacturer), as this cutting plane was perpendicular to the axis of the folded sample. After drying, the sections were stained with hematoxylin-eosin (BioVitrum, Russia). Additional staining with Alcian blue pH 3.0 was performed by contrasting the nuclei with Mayer’s haematoxylin, which allowed visualization of mucosal goblet cells. The samples were evaluated by light microscopy at X4, X10 and X40 magnifications (Zeiss Primo Star, China). A ball-based scoring system was used to assess the severity of damage, as previously described.

### HS-GC/MS

For HS-GC/MS fecal samples (all available sample volume) were collected every 3 days in the control, DSS and DSS+OMV groups. 50-100 mg stool samples plus 500 µL water samples were placed in 10 mL screw-capped vials for a Shimadzu HS-20 headspace extractor. 0.2 g of a salt mixture (ammonium sulfate and potassium dihydrogen phosphate in a ratio of 4:1) was added to increase the ionic strength of the solution. The following headspace extractor settings were used: oven temperature 80°C, sample line temperature 220°C, transfer line temperature 220°C, equilibration time 15 min, pressurization time 2 min, loading time 0.5 min, injection time 1 min, needle rinsing time 7 min. Vials were sealed and analyzed on a Shimadzu QP2010 Ultra GC / MS with a Shimadzu HS-20 headspace extractor, a VF-WAXMS column of 30 m length, 0.25 mm diameter and 0.25-micron phase thickness. Initial column temperature 80°C, heating rate 20°C/min to 240°C, exposure time 20 min. Carrier gas - Helium 99.9999, injection mode - splitless, flow rate 1 ml/min. Ion source temperature - 230°C. Interface temperature 240°C. Total ion current (TIC) monitoring mode used. The NIST 2014 Mass Spectra Library with Automated Mass Spectra Deconvolution and Identification System (AMDIS version 2.72) was used to analyzed the obtained mass spectra.

### Metabolome data processing

The HS-GC/MS data were processed as follows: The peak areas calculated by AMDIS for the selected compounds were converted into relative abundances. Volatile compound percentages were estimated by summing the percentages of confidently identified compounds for each sample in the AMDIS database. The resulting values were then recalculated as a percentage of the total number of compounds confidently identified. These conversions were necessary to avoid errors in content estimation due to unreliable matrix signals caused by noise.

The GC-MS data were processed using MetaboAnalyst 5.0 software (http://www.metaboanalyst.ca) and GraphPad Prism 8.0.1 software. The values obtained for each animal were considered paired and consistent, confirmed by the ROUT outlier test (ROUT (Q=1%)). As it was not possible to assess the normality of the distribution for all sample groups, it was assumed that the raw data did not conform to a normal distribution. The non-parametric Mann-Whitney test was used for the primary comparison between groups. After natural log transformation, normalized data were analyzed using a standard t-test and ANOVA. Statistical significance was determined by a two-tailed p-value of less than 0.05. Unsupervised principal component analysis (PCA) was used to reduce the number of dimensions and further explore the data. Standard procedures were applied prior to PCA. A chi-squared test was used to determine a method for imputing missing values. The missing at random (MAR) criteria were identified in the data. The BPCA method was found to be the most appropriate. Normalization and scaling procedures were then performed. Correlations between metabolites and gut microbiota composition were analyzed using the R ’psych’ package based on Spearman’s rank correlation coefficient [24].

### DNA extraction

Fecal samples were used for total DNA extraction. Nucleic acids were extracted using the MagicPure® Stool and Soil Genomic DNA Kit and Kingfisher Flex Purification System (Thermo Fisher Scientific, USA) according to the manufacturer’s protocol. DNA was then quantified using the Quant-iT dsDNA BR Assay Kit (Thermo Fisher Scientific, USA) on a Qubit 4 fluorometer.

### 16S rRNA gene sequencing on the MinION™ platform

The extracted DNA (1-5 ng) was amplified using 27F (AGAGTTTGATYMTGGCTCAG) and 1492R (GGTTACCTTGTTAYGACTT) primers (Eurogen, Russia) and the Tersus Plus PCR Kit (Eurogen, Russia) in a total volume of 25 µl. Amplification was performed with the following PCR conditions: initial denaturation at 95 °C for 2 min, (95 °C for 1 min, 60 °C for 1 min and 72 °C for 3 min), 27 cycles, followed by a final extension at 72 °C for 2 min and 4 °C - cooling. The quality of the amplicons was checked by electrophoresis in 1.5% agarose gel. The final amplicons were purified using KAPA HyperPure Beads (Roche, Switzerland) according to the manufacturer’s protocol.

Libraries were prepared according to the manufacturer’s protocol (ligation sequencing amplicons) with modifications. Amplicons were processed using the NEBNext® Ultra™ II End Repair/dA-Tailing Module (NEB). Barcodes (Native Barcoding Kit 96 (SQK-NBD109.96)) were ligated using Blunt/TA Ligase Master Mil (NEB). Barcoded libraries were purified using KAPA Pure Beads (Roche, Switzerland). Library concentrations were measured using the Quant-iT dsDNA Assay Kit, High Sensitivity (Thermo Fisher Scientific, USA) and samples were mixed at equimolar concentrations. The final adapter (Adapter Mix II Expansion (Oxford Nanopore Technologies, UK) was ligated to the pooled library using the NEBNext Quick Ligation Module (NEB). The prepared DNA library (12 μL) was mixed with 37.5 μL sequencing buffer, 25.5 μL loading beads, loaded onto the R9.4 flow cell (FLO-MIN106; Oxford Nanopore Technologies) and sequenced using the MinION™ Mk1B. MINKNOW software ver. 22.12.7 (Oxford Nanopore Technologies) was used for data acquisition.

### Genome data processing

Technical sequences and bases with a quality lower than a Phred score of 9 were processed using Porechop and NanoFilt software [24, 25]. The resulting data were evaluated using the Emu pipeline for taxonomic classification [26]. Alpha and beta diversity analyses were performed using the vegan package for GNU/R [27]. Alpha and beta diversity were assessed using the Bioconductor Microbiota Process package for GNU/R. Heatmap visualization was performed using the ’pheatmap’ package for GNU/R. Alpha diversity studies were performed using the Microbiota Process package [28, 29].

## Results

### Histological evaluation of intestine inflammation and OMV therapy

Isolated OMVs were visually characterized by TEM. As previously described, vesicles were approximately spherical and most particles had a relatively dark part in the center (**Figure S1**) [15]. The OMVs obtained were used to treat mice after DSS exposure. All three groups were subjected to histological examination of intestinal tissues on the 10^th^ and 20^th^ day of the experiment. According to the histopathological results (**Fig. 1А**), the mucosa and submucosa of the colon in the control group (**Fig. 1А K1-K2**) were thin without signs of oedema. The mucosa did not have extensive areas of damage along its entire length and retained smooth, slit-like crypts. Blood and lymphatic vessels were not dilated. The epithelium with mild inflammatory changes in several animals from the control group was characterized by non-extended areas of goblet cell loss (**Figs 1A K3**). In the DSS group (**Fig 1A DSS 1-2**), the structure of the mucosa in the distal parts of the colon was not determined; only individual goblet cells were observed. (**Fig. 1A DSS 3**). In other parts of the colon, focal lesions of the mucosa were seen to a lesser extent. The lesions showed loss of normal epithelial structure, dilated, branched crypts of reduced depth and abundant immune cell infiltration. Crypt abscesses and foci of mucosal infiltration were identified in the proximal intestine. An accumulation of immune cells was observed in the submucosal layer of the colon of all experimental animals. The mucosa and submucosa were slightly thickened, and blood and lymphatic vessels were dilated. A smaller lesion area was observed in the DSS OMV group compared to the DSS group (**Fig. 1A DSS OMV 1-2**). Mucosal foci were also identified in the distal sections of the colon, but their structure was indistinguishable and the number of goblet cells was reduced (**Fig. 1A DSS OMV 3**). However, in contrast to the DSS group, the lesion was focal in nature and had a significantly shorter extent. In the distal parts of the bowel, the lesions occurred over a longer period of time and were also more pronounced. Crypt abscesses and areas of ulceration were also seen in both the distal and proximal bowel, but only in a few cases than in the DSS group. In areas of mucosal lesions, crypts were dilated and the number of goblet cells was also reduced. Many nodules of lymphoid tissue were seen in the submucosal layer. The mucous and submucous membranes were slightly thickened and the blood and lymphatic vessels were dilated. Thus, histological examination of the bowel samples from the control group showed no signs of a massive inflammatory process. The characteristic histological signs of colitis were not seen. In the DSS group, a characteristic picture of chronic colitis was observed, including changes in the epithelium (increased thickness of the submucosal and mucosal layers of the bowel, expansion or complete disappearance of crypts, inflammatory infiltration of the mucosa and submucosal layer, decreased number of goblet cells, increased number of lymph nodes in the submucosal layer). In the DSS OMV group, characteristic histological signs of chronic colitis were also observed, including focal changes in the epithelium, but the extent and degree of manifestation were less than in the DSS group, suggesting a reduction in disease severity. The histopathological scoring system was used to detect epithelial damage and inflammatory cell infiltration. According to the data obtained (**Supplementary Table 1**), the histological index was significantly lower in the DSS OMV group than in the DSS group, indicating tissue repair under OMV treatment (**Fig. 1B**). As shown in **Figure 1С**, the DAIs of the DSS and DSS OMV groups were significantly higher than those of the control group.

**Figure 1.**
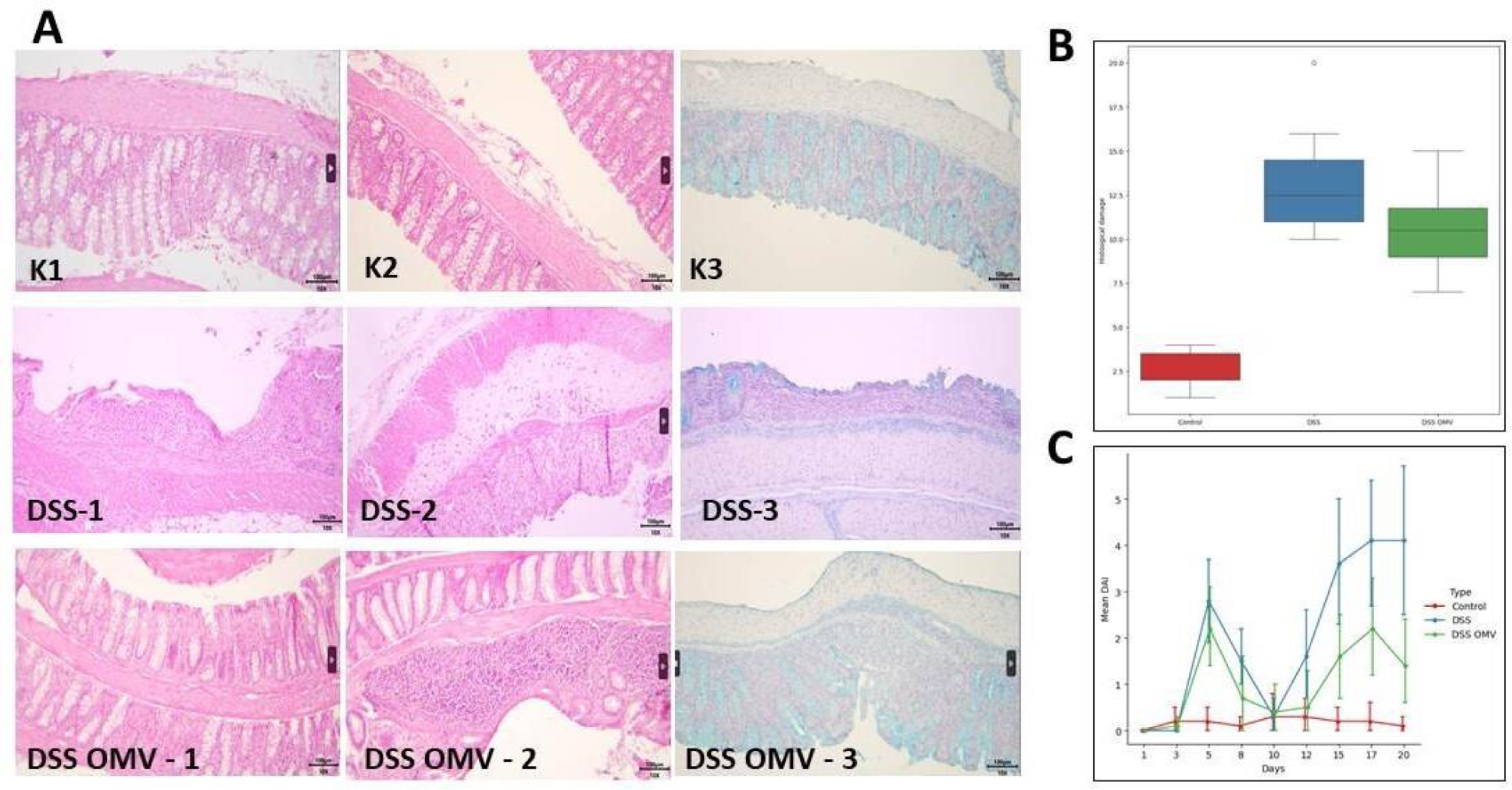
Histological examination of intestinal tissue for all groups. A - Formalin-fixed preparation of the distal intestine of an experimental animal. Hematoxylin-eosin staining. (K1,K2,DSS1,DSS2,DSSOMV-1,DSSOMV-2). Additional staining with Alcian blue pH 3.0 was performed by contrasting the nuclei with Mayer’s hematoxylin, which allowed visualization of mucosal goblet cells (K3, DSS3, DSSOMV-3). K1, K2, K3 - control (without DSS); DSS-1, DSS-2, DSS-3 - DSS-treated intestine; DSSOMV - 1, DSSOMV - 2, DSSOMV - 3 - OMV-treated intestine. The samples were evaluated by light microscopy at X4, X10 and X40 magnifications (Zeiss Primo Star, China). B - Evaluation of histological sections of the distal intestine in points. C - DAI for all three groups throughout the experiment, day by day.

DAIs in both groups increased until day 3, mainly due to weight loss of the animals, and peaked on day 5, when severe diarrhea with visible blood in the stool was observed. DAIs decreased in both groups by day 10 after discontinuation of DSS. When DSS was resumed, an increased DAI was again observed in both groups. However, when OMVs therapy has been implicated a significant decrease in the growth rate of DAI in the DSS OMV group compared to the DSS group (**Supplementary Table 1**).

### Metabolomic data

The metabolite spectrum was evaluated for samples collected on the 1^th^, 10 ^th^ and 20 ^th^ day of the experiment. The study was performed by HS-GC/MS. Metabolomic data was normalized (**Supplementary Table 2**). At least 60% of the detected stable compounds were used for the metabolomic profiles comparison. The most frequently detected compounds meeting the screening criteria are listed in **Table 1**.

**Table 1.** Complete list of stably detected metabolites in three checkpoints days of the experiment.

| Metabolite | Percent of samples % |  |  |  |  |  |  |  |  |  |
| --- | --- | --- | --- | --- | --- | --- | --- | --- | --- | --- |
|  | Control group (d) |  |  | DSS group (d) |  |  | DSS+OMV group (d) |  |  | Percent of samples in which the metabolite is detected<br>n=263 |
|  | 1 | 10 | 20 | 1 | 10 | 20 | 1 | 10 | 20 |  |
| Acetic acid | 90.9 | 90.9 | 90.9 | 90.9 | 90.9 | 90.9 | 90.9 | 90.0 | 90.9 | 100.0 |
| Propanoic acid | 90.9 | 90.9 | 90.9 | 90.9 | 90.9 | 90.9 | 90.9 | 90.0 | 90.9 | 100.0 |
| Benzaldehyde | 90.9 | 72.7 | 90.9 | 72.7 | 72.7 | 72.7 | 72.7 | 50.0 | 63.6 | 79.8 |
| Propanoic acid, 2-methyl- | 90.9 | 90.9 | 90.9 | 90.9 | 90.9 | 90.9 | 90.9 | 90.0 | 90.9 | 100.0 |
| Benzeneacetaldehyde | 90.9 | 72.7 | 81.8 | 90.9 | 63.6 | 90.9 | 90.9 | 50.0 | 72.7 | 87.8 |
| Butanoic acid, 3-methyl- | 90.9 | 90.9 | 90.9 | 90.9 | 90.9 | 90.9 | 90.9 | 80.0 | 90.9 | 99.6 |
| Methyl formate | 63.6 | 81.8 | 72.7 | 81.8 | 63.6 | 54.5 | 90.9 | 50.0 | 81.8 | 72.2 |
| Butanoic acid, 2-methyl- | 90.9 | 90.9 | 90.9 | 90.9 | 81.8 | 81.8 | 90.9 | 90.0 | 90.9 | 96.6 |
| Pentanoic acid | 90.9 | 90.9 | 90.9 | 90.9 | 90.9 | 90.9 | 90.9 | 90.0 | 90.9 | 100.0 |
| Pentanoic acid, 4-methyl- | 72.7 | 90.9 | 81.8 | 90.9 | 54.5 | 72.7 | 90.9 | 20.0 | 45.5 | 77.6 |
| Phenol | 90.9 | 90.9 | 90.9 | 90.9 | 72.7 | 72.7 | 90.9 | 40.0 | 90.9 | 81.7 |
| Octanoic Acid | 9.1 | 90.9 | 81.8 | 90.9 | 63.6 | 63.6 | 90.9 | 50.0 | 81.8 | 79.1 |
| Benzyl Alcohol | 90.9 | 90.9 | 90.9 | 90.9 | 81.8 | 81.8 | 90.9 | 90.0 | 90.9 | 97.3 |
| Undecanal | 72.7 | 72.7 | 54.5 | 63.6 | 63.6 | 27.3 | 81.8 | 50.0 | 72.7 | 76.4 |
| Nonanoic acid | 63.6 | 90.9 | 90.9 | 90.9 | 45.5 | 90.9 | 90.9 | 80.0 | 81.8 | 89.4 |
| 5-Acetyl-2-methylpyridin | 81.8 | 72.7 | 54.5 | 54.5 | 72.7 | 27.3 | 63.6 | 70.0 | 54.5 | 67.7 |
| Indole | 90.9 | 90.9 | 81.8 | 90.9 | 81.8 | 72.7 | 72.7 | 80.0 | 72.7 | 90.9 |
| Tetradecanal | 90.9 | 72.7 | 54.5 | 81.8 | 9.1 | 63.6 | 72.7 | 40.0 | 45.5 | 64.6 |
| Benzenepropanoic acid | 36.4 | 63.6 | 81.8 | 72.7 | 9.1 | 81.8 | 90.9 | 30.0 | 90.9 | 63.5 |
| Heptanoic acid | 18.2 | 72.7 | 90.9 | 90.9 | 45.5 | 63.6 | 90.9 | 30.0 | 81.8 | 71.9 |
| Phenylethyl Alcohol | 81.8 | 63.6 | 81.8 | 81.8 | 45.5 | 45.5 | 81.8 | 70.0 | 90.9 | 70.7 |

Among the compounds identified, SCFAs, medium and long chain fatty acids and amino acid derivatives were detected. Differences in the metabolic profiles of the control group and the DSS and DSS OMV groups were shown using PCA and OPLS DA (**Fig. 2 A, B**). According the data obtained samples were clustered into three independent groups. The control group was distinguished by a separate characteristic spectrum of the relative abundance of volatile compounds in the DSS and DSS OMV groups on the 10^th^ day of the experiment. No significant differences were found between the DSS and DSS OMV groups on day 10 of the experiment. On day 20, corresponding to the end of the vesicle’s treatment, three separate groups could be distinguished. The control group was significantly different from the DSS and DSS OMV groups. A small number of metabolomic differences could be identified between the DSS and DSS OMV groups according to the PCA data. OPLS DA allowed the assessment of the magnitude of the differences, which contributed to a clearer classification of the DSS and DSS OMV groups.

**Figure 2.**
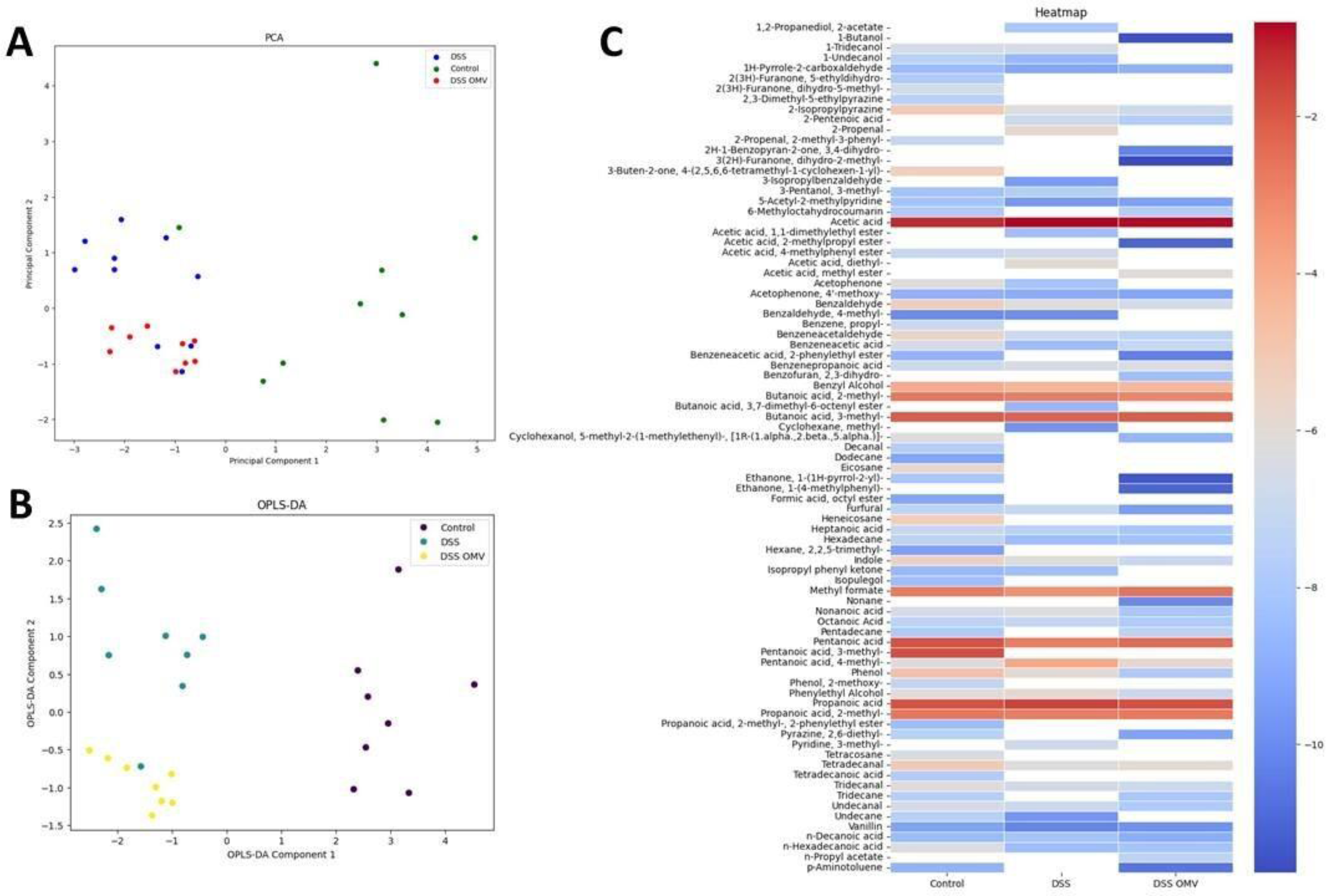
Total HS-GC/MS data for all groups on the 20th day of the experiment. **A, B** - PCA and OPLS DA data represent three independent groups of samples, according to the relative concentration of volatile compounds. **C** - VOCs compounds identified in all groups. The relative concentrations in the vapor phase are used.

General profile of DSS OMV group was characterized by 2,5-Dimethyl-4-hydroxy-3(2H)- furanone, H-1-benzopyran-2-one, 3,4-dihydro, nonane which were not detected in the control and DSS groups (**Fig. 2C**). The overall profile of metabolites was significantly depleted in the DSS group compared to the control and DSS OMV groups. When treated with vesicles, a tendency to restore the overall metabolite profile could be observed (**Fig. 2C**).

Significant changes in the relative amounts of metabolites were observed both when comparing the control and DSS OMV groups and when comparing the DSS and DSS OMV groups on day 20. The main changes were affected such compounds as acetic acid, propanoic acid, propanoic acid 2-methyl, benzaldehyde, pentanoic acid, phenol, nonanoic acid, heptanoic acid and hexanoic acid (**Fig. 3 A, B**). Mentioned metabolomic differences were identified in all the three groups on the 20th day (**Figure 3 C)**.

**Figure 3.**
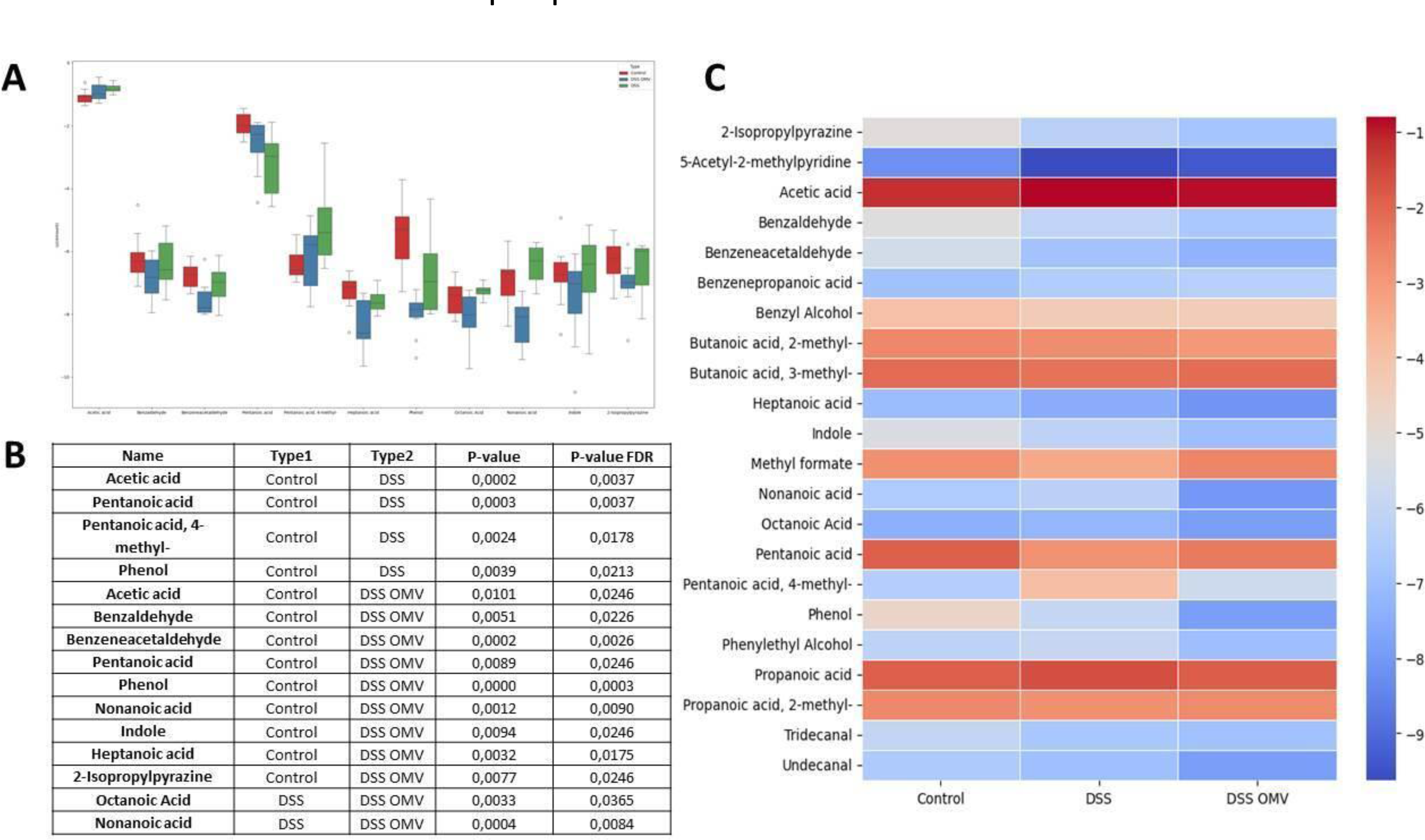
Quantitative differences of individual components in experimental groups. **A** - Box plots show the quantitative differences in the relative content of short-chain fatty acids, amino acid derivatives and others detected in the analyzed groups. **B** - The non-parametric Mann-Whitney test was used for the primary comparison between groups. Statistical significance was determined by a two-sided p-value of less than 0.05. FDR correction was also applied. **C** - Comparison of VOC composition on day 20 in all groups. Relative concentrations in the vapor phase are used.

Most of the significant differences were found between the control and DSS OMV groups on day 20. It can be assumed that after OMVs treatment a new equilibrium of the relative amounts of individual metabolites can be observed.

### Amplicon sequencing data analysis

Sequencing analysis of microbiome variability in stool samples was performed on the 1^th^, 10 ^th^ and 20 ^th^ day of the experiment (**Supplementary Table 3**). As shown in **Figure 4** (**Supplementary Figure S2**), at the beginning of the experiment a homogeneity of bacterial species was observed in the analyzed groups. At the same time, there was a significant predominance of *Fermicutes* in the microbiota. On the tenth day of the experiment, a decrease in number of several bacterial species was observed in the DSS and DSS OMV groups compared to the control. As expected, uniformity was still observed in the DSS and DSS OMV groups on day 10. However, by day 20 there was a significant difference between the DSS and control groups. It can be seen that there was a significant reduction in the number of bacterial species in the DSS group compared to the control group. However, in the DSS OMV group receiving vesicles, a greater diversity of bacterial species could be observed at the end of the experiment. The numerical bacterial diversity in the DSS OMV group approaches the initial diversity observed in the control group. Tables showing the relative abundance of bacterial species in each sample and statistics of processed reads can be found in the **Supplementary table 3**.

**Figure 4.**
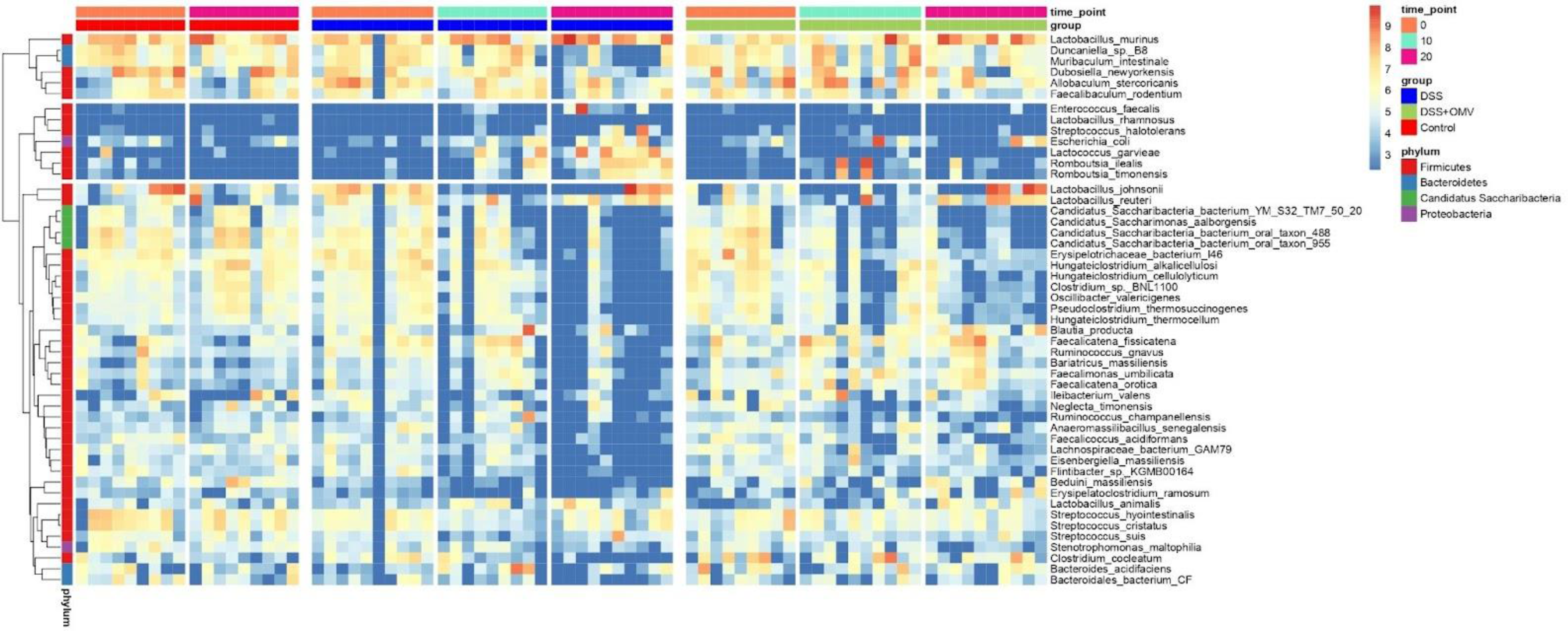
Bacterial diversity in different time points (1,10,20 days). The color scale of the heat map shows the abundance of different species. The horizontal axis represents samples, the vertical axis represents different bacterial species.

Analysis of the relative abundance of bacteria at the genus level allowed us to follow the change in bacterial composition from the control group to the DSS+OMV group. According to the data obtained, the relative abundance within the studied bacterial genera in the DSS+OMV group reached the initial values determined in the control group by the twentieth day of observation. In contrast to the DSS group, where dramatic changes occur, a tendency for the number of microorganisms in the DSS+OMV group to return to control values could be observed when the vesicles were used (**Fig. 5A**). A Venn diagram and upset plot can be used to visualize the overlap in bacterial species between the three groups shown. The DSS+OMV group was close in species composition to the control group (117 species). The DSS OMV group and the DSS group had 30 species in common. The DSS OMV group had more than 100 bacterial species in common with the control group, whereas the DSS group and the control group had only 30 species in common (**Fig. 5 B,C**).

**Figure 5.**
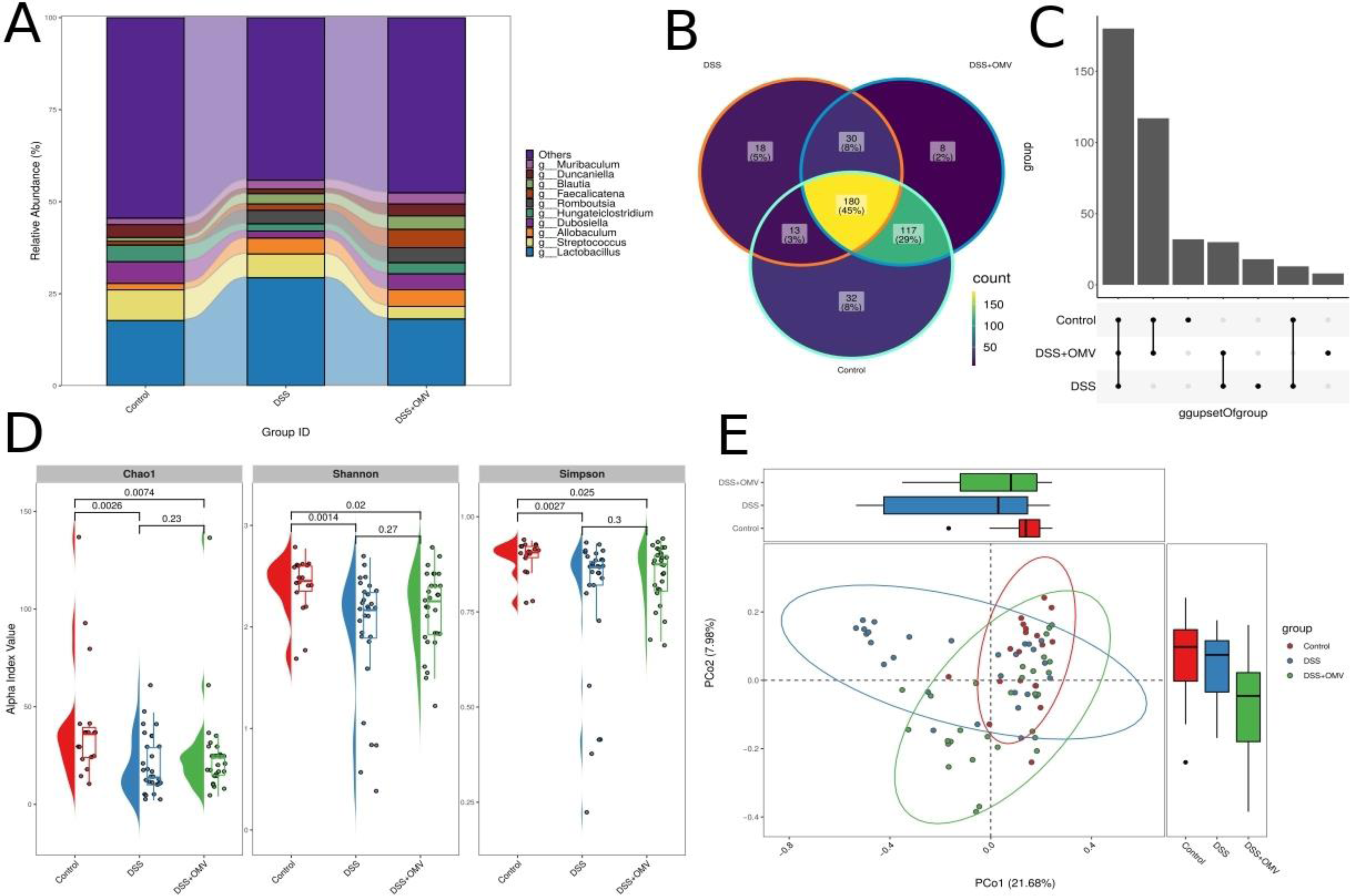
Bacterial diversity in different time points (1,10,20 days) A - Relative abundance of bacteria at genus level for each group (control, DSS, DSS+OMV) across all sites. The horizontal axis shows groups and the vertical axis shows relative abundance. B,C - Venn diagram and upset plot for groups (control, DSS, DSS+OMV) at OTU level. D - Plot of alpha diversity index at all points. The horizontal axis represents each group (control, DSS, DSS+OMV), the vertical axis represents the alpha diversity index. E- PCoA plot based on Bray-Curtis distance for all time points. Each point represents one sample. The color of the point indicates the name of the sample group.

An alpha diversity study was performed to assess the homogeneity and richness of the microbial community composition. The result of the analysis showed that the alpha diversity value for the three groups was statistically significantly different only on day 20. Species diversity in the DSS group was significantly different from the control and vesicle-treated groups on day 20 of the experiment. When assessing alpha diversity, it could be seen that all three groups were significantly different on the last day of the experiment, but the degree of difference between the control group and the DSS group was more pronounced than the differences between the control group and the group receiving vesicles (DSS OMV) (**Fig. 5 D**). Beta diversity analysis was used to quantify the differences between the bacterial communities of the samples. Principal Coordinate Analysis (PCoA) showed that the bacterial communities were statistically different between all three groups (**Fig 5 E**). The Manhattan plot also showed significant diversity between three groups (**Figure S3**). LDA analysis was performed to search for significant bacterial taxa for each experimental group [28]. The result showed that the OTUs were significantly enriched in the control group and were belonged mainly to the phylum *Candidatus Saccharibacteria* and the genus *Hungatei Clostridium*. The OTUs which were significantly enriched in the DSS group were mainly from the genera *Lactococcus* and *Romboutsia* but the OTUs which were significantly enriched in the DSS+OMV group were mainly from the genus *Faecalimonas* (**Figure 6**).

**Figure 6.**
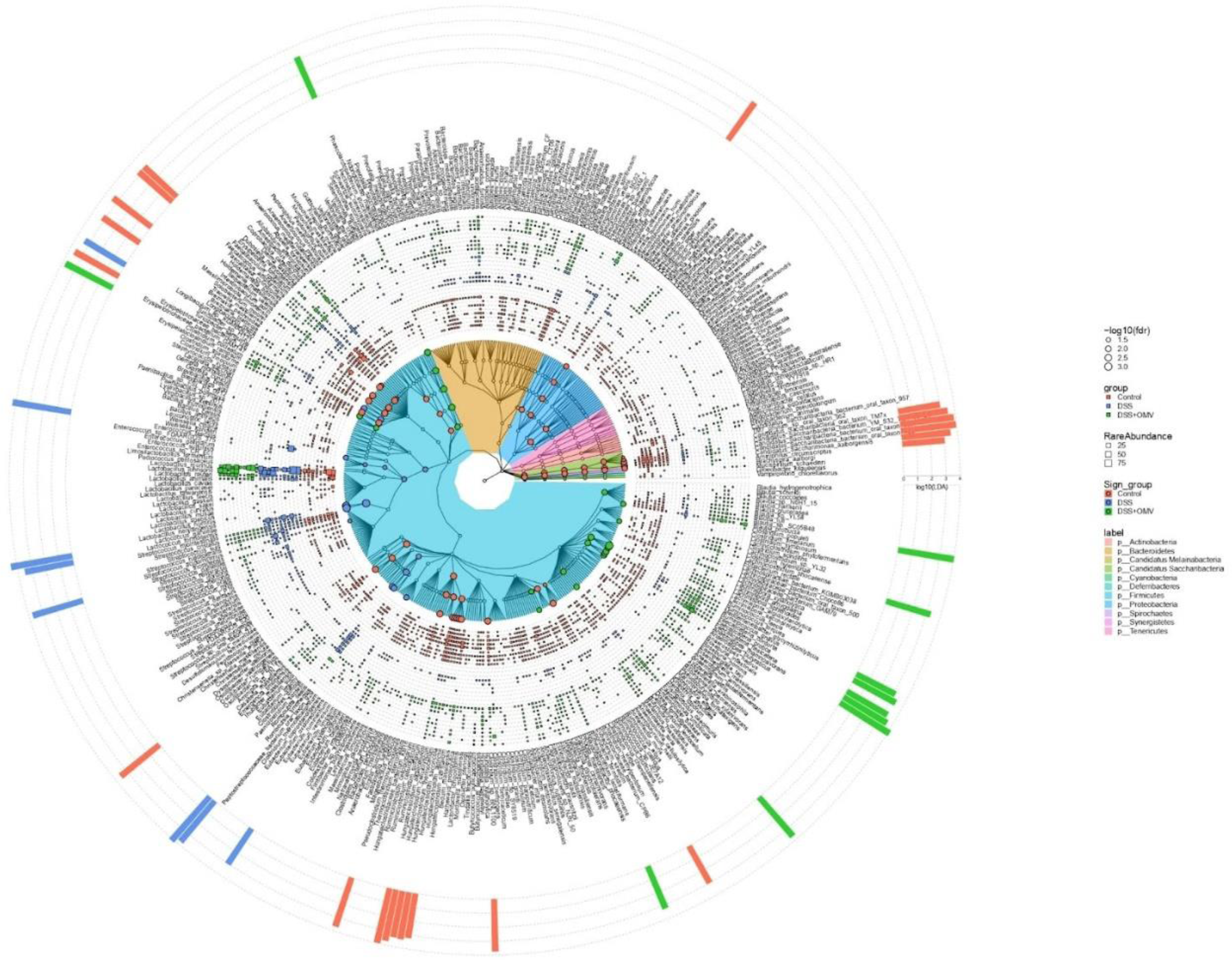
Taxonomic community tree with relative abundance of each OTU in the sample and LDA of different OTUs. The tree was constructed using taxa from all samples. Coloured clades represent phyla. The outer layer shows the relative abundance of each OTU in the sample. The outer histogram shows the LDA of different OTUs. The coloured dots represent different taxa and the dot size indicates the FDR.

Different correlations between microbiota composition and metabolomic compounds such as phenol and pentanoic acid were found **Figure 7**. When comparing the relative amounts of these metabolites and the diversity indices, a significant correlation could also be observed. According to the data obtained, in the control group there was a predominance of *Saccharibacteria* and *Hungatei clostridium* were positively correlated with phenol and pentanoic acid and the DSS group there was a predominance of *Lactococcus* and *Romboutsia were* negatively correlated with phenol and pentanoic acid.

**Figure 7.**
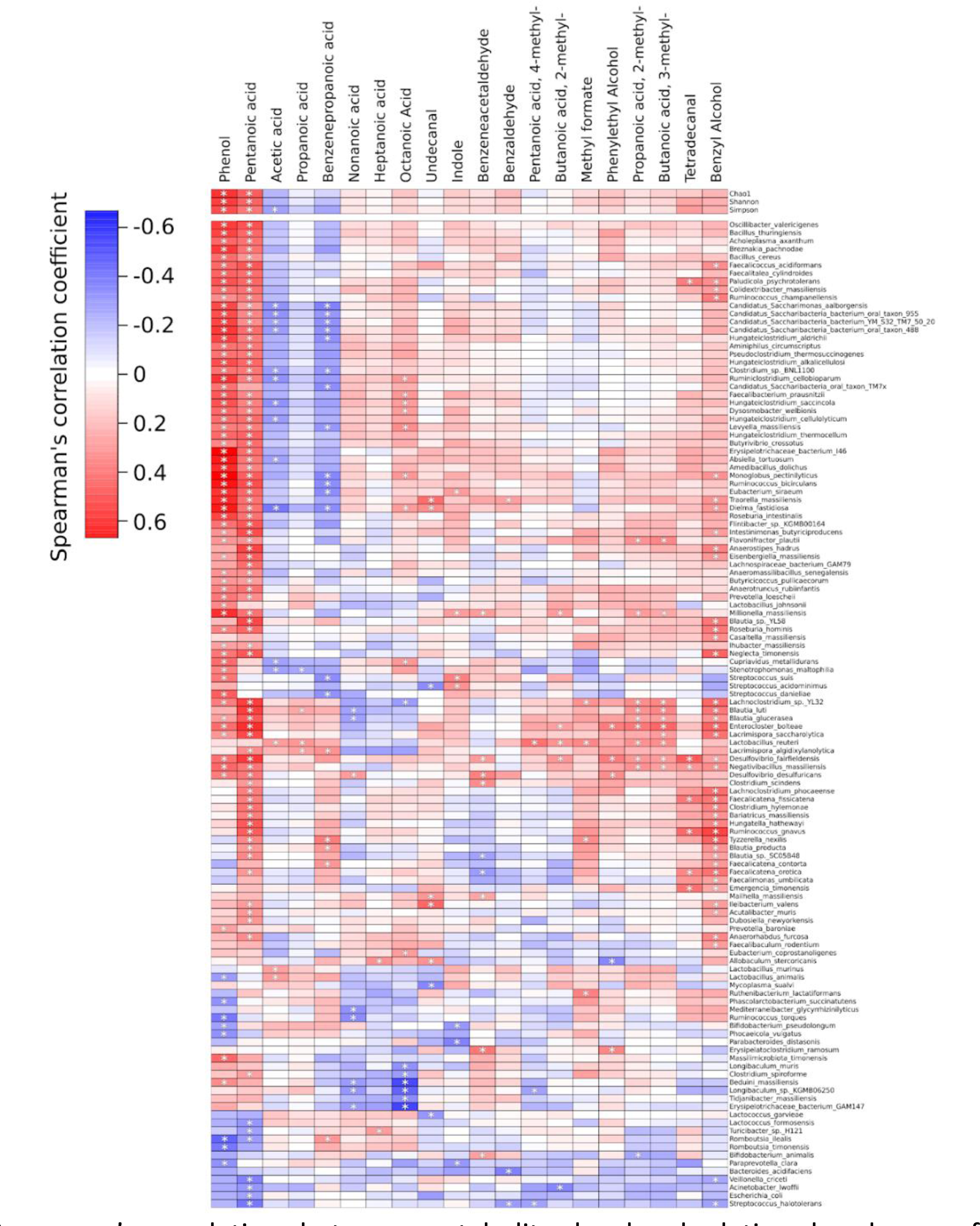
Spearman’s correlations between metabolites level and relative abundances of microbial species. * - p <0.05

## Discussion

DSS-induced colitis is a useful experimental model for studying the pathogenesis of inflammatory bowel disease and for testing new therapeutic agents [30,31]. DSS causes intestinal damage and triggers the formation of all stages of classical inflammation [32]. The immune system is also involved in this process, which is of paramount importance in inflammatory bowel diseases such as Crohn’s disease and ulcerative colitis due to the development of autoimmunity [33]. Therefore, existing therapies are mainly aimed at suppressing the autoimmune response that provokes chronic intestinal inflammation [34]. With the development of a chronic inflammatory process, the microbiota inevitably suffers [35]. It has been shown that with the development of Crohn’s disease and ulcerative colitis, a different ratio of bacterial species is formed, as well as the appearance of a greater number of pathogenic bacteria [36]. The morphological changes lead to a decrease in the activity of the symbiotic microbiota, which has a negative effect on the gut [37]. In addition, the normal microbiota contains unique species whose influence on the intestinal mucosa helps to reduce the inflammatory response [38]. *Bacteroides fragilis* secretes vesicles with polysaccharide A (PSA) on their surface [8]. Some studies have shown that isolated PSA improves mucosal recovery in an induced colitis model [16]. The use of OMVs for the treatment of IBD is more justified than the use of isolated PSA because vesicles contain a significant number of enzymes that help to improve digestive function [14]. In this study, we recreated a model of intestine inflammation in the presence of DSS and treated it with *Bacteroides fragilis* JIM10 vesicles for ten days. We first focused on the histological and physical assessment of the experimental animals. As expected, on day 10 of DSS exposure we observed changes in the intestinal tissue according histopathology examination. However, as early as day 20 of the experiment, when we used OMVs, we observed a positive therapeutic effect, which was approved by histological examination. Compared to the group without OMVs treatment, the group that received vesicles was better able to repair the structure of the intestinal crypts and reduce the amount of inflammatory infiltrate in the tissue. In fact, we obtained an effect similar to that previously demonstrated with isolated PSA [16]. When we analyzed the microbiota composition, we observed bacterial diversity in all three groups. For alpha and beta diversity, we observed changes in both the DSS and DSS-OMV groups. It is important when OMVs were used as therapy, there was a tendency for the microbiota composition in DSS OMV group to return to its original control abundance. However, each group was characterized by its own unique bacterial diversity on day 20. Our results showed that *Candidatus saccharibacteria* and the genus *Hungatei* were mainly enriched in the control group. *Lactococcus* and *Romboutsia* were significantly enriched in the DSS group and the genus *Faecalimonas* was significantly enriched in the DSS+OMV group. It can be assumed that the OMVs contributed to the recovery of the original gut homeostasis which is combined health mucosa layer and specific microbiota composition. Nevertheless, the changes in microbiota composition under DSS treatment were not as dramatic as might be expected. However, the demonstrated OMVs influence on microbiota composition is unique.

Metabolomic data has complemented 16S rRNA sequencing data. Changes in relative amounts of volatile components, such as short-chain fatty acids, phenol and pentanoic acid, were detected. Correlation of metabolites such as phenol and pentanoic acid with indices of microbiota diversity in the groups was observed. Relative amounts of mentioned metabolites were decreased in both experimental groups. *Saccharibacteria* and *Hungatei сlostridium* were positively correlated with phenol and pentanoic acid in the control group, whereas *Lactococcus* and *Romboutsia* were negatively correlated with phenol and pentanoic acid in the DSS group. Based on the metabolomic and genome data obtained, we can clearly conclude that DSS can induce changes in the microbiota composition, but *Bacteroides fragilis* OMVs contribute to the recovery of the original ratios.

We hypothesize that vesicles can either directly influence the microbiota composition delivering enzymes and metabolites necessary to restore microbiota functional activity. On the other hand, the intestinal mucous layer repair under OMVs treatment can also contribute to the recovery of the original microbiota composition.

## Conclusion

The aim of the present study was to evaluate the therapeutic effects of Bacteroides fragilis JIM10 OMVs in recovery process of DSS-affected intestinal tissue in murine model. According to the data obtained, it was possible to demonstrate the therapeutic effectiveness of Bacteroides fragilis JIM10 OMVs and also to observe the unique OMVs property to contribute the recovery of intestinal microbiota composition.

## Authors contribution

O.Y.S., D.A.K.,V.A.I., E.A.V., M.I.M., E.I.O., N.B.Z. designed and performed experiments, analyzed data and wrote the paper; D.A.K.,S.S.E.,E.A.Z.,O.V.A., Ya.A.Z., A.Y.M., A.S.S., B.A.E., A.A.V., M.D.M., O.Y.Z., D.I.B. performed experiments; T.V.G., N.B.Z.supervised the project.

## Acknowledgements

The TEM measurements were carried out at the User Facilities Center “Electron microscopy in life sciences” of Lomonosov Moscow State University. We thank Mr. Alexey Senkovenko and Mr. Dmitry V. Bagrov for the assistance with TEM imaging.

